# Circulating microRNA-762 upregulates colorectal cancer might through Wnt-1/β-catenin signaling

**DOI:** 10.1101/2020.04.22.056796

**Authors:** Peng-Sheng Lai, Wei-Min Chang, Ying-Yin Chen, Yi-Feng Lin, Hui-Fen Liao, Chung-Yu Chen

## Abstract

**Background:** A number of microRNAs (miRNAs) have been demonstrated to be correlated with the diagnosis, progression and prognosis of colorectal cancer (CRC). However, the key miRNAs and the associated signaling pathways that regulate the growth and metastasis of CRC remain unclear.

**Methods:** The circulating miRNAs from BALB/c mice with CRC CT26 cell implantation were assayed by microarray. Then, mmu-miR-762 mimic and inhibitor were transfected to CT26 cells for analysis of cell viability, invasion, and epithelial-mesenchymal transition (EMT), cell cycle, and regulatory molecule expression. Human subjects were included for comparison the circulating has-miR-762 levels in CRC patients and control donors, as well as the patients with and without distant metastasis.

**Results:** The miRNA levels in mice with CRC cell implantation indicated that plasma mmu-miR-762 was upregulated. Transfection of mmu-miR-762 mimic to CT26 cells increased cell viability, invasion, and EMT, whereas transfection of mmu-miR-762 inhibitor decreased the above abilities. Cells treated with high-concentration mmu-miR-762 inhibitor induced cell cycle arrest at G0/G1 phase. Western blot analysis showed that mmu-miR-762 mimic transfection upregulated the expression of Wnt-1 and β-catenin. Further analysis was performed to demonstrate the correlation of has-miR-762 with CRC patients. The results showed that serum has-miR-762 levels in CRC patients were higher than in control donors. Among the CRC patients, patients with distant metastasis showed higher serum has-miR-762 levels than patients without distant metastasis.

**Conclusions:** The present study demonstrated that circulating miR-762 might be a biomarker with upregulation of CRC cell growth and invasion through the Wnt/β-catenin signaling.

## 1. Introduction

Colorectal cancer (CRC) is epithelial in origin and localized to the large intestine and rectum. CRC has been ranked among the top five most common cancers worldwide. Because of the aging population, westernization of diet, excessive intake of fat, reduced intake of dietary fiber, and other factors, the incidence of CRC continues to increase and has become more prevalent in younger people^1^. The most common treatment for CRC is surgery to remove the tumor before metastasis occurs, which shows a very high cure rate. Moreover, combinations of chemotherapy drugs such as 5-fluorouracil, camptothecin, bevacizumab, and cetuximab are effective for CRC treatment^2^. However, metastasis is the cause of death in most CRC cases; depending on the tumor stage, liver and lung metastases occur in 20–70% and 10–20% of cases, respectively^3^. It has been reported that approximately 50% of patients with CRC develop tumor metastases which are associated with very poor prognosis and treatment efficiency^4^.

MicroRNAs (miRNAs) are non-translated RNAs that regulate gene expression by degrading mRNA or inhibiting translation^5^. This endogenous mechanism is common to animals, plants, and viruses. miRNAs have been shown to be involved in many biological processes, such as embryonic development, inflammation, cell cycle regulation, cell differentiation, apoptosis, and tumor metastasis^6^. In recent years, studies have shown that the occurrence of cancer and tumor formation are closely related to miRNA regulation of transcriptional gene expression^7^ and miRNAs can be used as indicators for diagnosing cancer^8^. For example, the miRNAs involved in the development of CRC and other cancers include miR-17-5p^9^, miR-21^10^, miR-31^11^, miR-92^12^, miR-198^13^, and miR-203^14^. These miRNAs can be used as biomarkers for cancer diagnosis diagnose or targeted therapy.

Therefore, identifying miRNAs that regulate the growth of CRC and clarifying the regulation of tumor formation and metastasis may reveal disease-specific biomarkers to detect the occurrence or prognosis of cancers. These results would also provide an important foundation for the research and development of advanced treatments in new target therapy strategies. In our previous miRNA array analysis in animal experiments, we detected some miRNAs that may be involved in the growth of CRCs. Among these, plasma mmu-miR-762 was found to be upregulated in CRC CT26 cell-transplanted BALB/c mice, suggesting that miR-762 is a potential biomarker for CRC growth regulation. Therefore, we assessed the effects of mmu-miR-762 on the viability, colony formation, invasion, cell cycle, and regulatory molecules of CRC CT26 cells by transfection with miRNA mimics and inhibitors. Finally, blood samples were collected from patients with CRC, and the effects of has-miR-762 in control subjects and cancer cases were compared. These results by analysis in cell line, mouse, and human subjects would provide insight into the role of miR-762 in regulating CRC.

## 2. Materials and methods

### 2.1. Cells and cell culture

CT26 cells are undifferentiated CRC cell lines produced in N-nitroso-N-methylurethane-induced BALB/c mice. The cells were purchased from American Type Culture Collection (ATCC, Manassas, VA, USA). CT26 cells were cultured in RPMI-1640 medium (Gibco, Grand Island, NY, USA) supplemented with 2 mM L-glutamine and 10% heat-inactivated fetal bovine serum (FBS; Gibco) at 37°C, passaged every 2–3 days with TEG solution (0.25% trypsin, 0.1% EDTA, and 0.05% glucose in Hanks’ balanced salt solution), and maintained at exponential growth.

### 2.2 Animal experiments and mouse miRNA arrays

BALB/c male mice (6–8 weeks old), obtained from the National Laboratory Animal Center (Taipei, Taiwan), were maintained in a pathogen-free environment and allowed free access to food and water. All animal experiments were performed according to the guidelines for the care and use of research animals (DHHS publ. NIH 85-23, revised 1996) and the animal use protocol had been reviewed and approved by the institute animal care and use committee (IACUC) of National Yang-Ming University, Taipei, Taiwan (Approval no. 100609). The mice were divided into two groups: normal control (n = 3) and CRC CT26 cell implantation (n = 3). As shown in **Figure 1A**, CT26 cells were prepared as a single-cell suspension in sterile phosphate-buffered saline (PBS) and injected subcutaneously (*s.c.*) with 2 × 10^5^ cells in the back of mice. Fourteen days after tumor implantation, the tumors had grown to a diameter of 3–5 mm, and then the mice were sacrificed by cervical dislocation euthanasia and blood samples were collected from each mouse. Plasma was analyzed by miRNA array (Phalanx Biotech Group, Inc., Hsinchu, Taiwan) according to the manufacturer’s protocol. Clustering was performed to visualize the correlations among replicates and under varying sample conditions. Up- and down-regulated genes are represented in red and green colors, respectively. Standard selection criteria to identify differentially expressed miRNAs were established as log2|fold-changeļ > 0.8 and P < 0.05.

**Figure 1.**
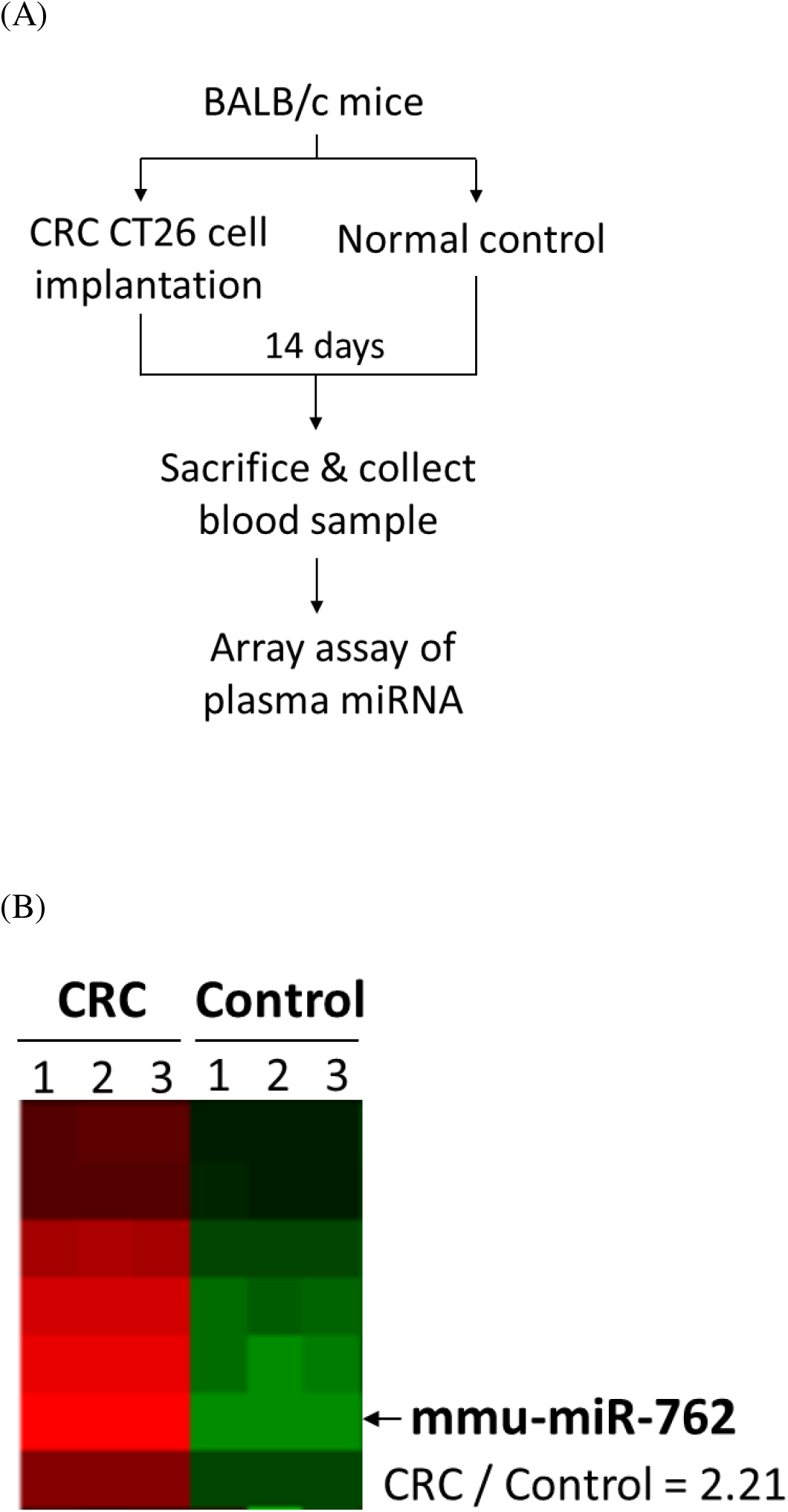
Array assay of plasma miRNA in BALB/c mice with CT26 cell implantation. (A) Animal experiment process, (B) Plasma mmu-miR-762 expression by miRNA array. Clustering was performed to visualize the correlations among the replicates and varying sample conditions. Up- and down-regulated genes are presented in red and green colors, respectively.

### 2.3 Transfection of mmu-miR-762 mimic and inhibitor

CT26 cells were cultured in a petri dish for 4 h before transfection. miR-762 mimic and inhibitor (miR-Ribo^TM^ miRNA, RiboBio Co., Guangzhou, China) were diluted with serum-free RPMI-1640 medium to final concentrations of 50 and 150 nM, mixed well, and incubated at room temperature (25 ± 2 °C) for 5 min. Next, the mmu-miR-762 mimic and inhibitor were mixed with Lipofectamine^TM^ 2000 transfection reagent (Invitrogen, Carlsbad, CA, USA), incubated for 20 min, and added to cultured CT26 cells. After 6 h, the medium was removed and fresh RPMI-1640 medium supplemented of 2 mM L-glutamine and 10% FBS was added for further culture for 24 h.

### 2.4 Cell viability assay

Cultured CT26 cells were collected and their viability was assayed by the 3-(4,5-dimethylthiazol-2-yl)-2,5-diphenyltetrazolium bromide (MTT, Sigma, St. Louis, MO, USA) colorimetric method. Briefly, treated cells were collected, washed with PBS, and reacted with MTT reagent (Sigma) for 2 h at 37°C to allow conversion of tetrazolium salts to a colored formazan product. The optical density of each reaction was measured spectrophotometrically at 550 nm to calculate cell viability. Cell morphology was observed under an inverted microscope (Olympus, Tokyo, Japan) at a magnification of 200X.

### 2.5 Colony formation assay

Viable cells (10^3^) were plated into 35-mm culture dishes and allowed to grow in RPMI-1640 medium containing 20% heat-inactivated FBS and 0.3% agarose at 37°C in a humidified 5% CO_2_ incubator. After 12 days, the dishes were stained with 0.4% crystal violet and colony numbers (1 colony contained more than 50 cells) were counted.

### 2.6 Cell invasion assay

Assays of cell invasion properties were performed using a modified Boyden chamber with polyethylene terephthalate filter inserts coated with a Matrigel matrix in 24-well plates containing 8-mm pores. Briefly, the cells were suspended in serum-free medium containing 0.5% bovine serum albumin. These cells were plated into the upper chamber followed by filling the lower chamber with the same medium with or without miR-762 mimic and inhibitor. Cells were incubated for 24 h, and then non-invading cells were gently removed. Cells on the upper side of the filter were carefully removed and cells invading the lower side were counted by microscopic examination (100X, Olympus). For quantification, 10% acetic acid (100 μL/well) was used to dissolve stained cells, which were transferred to 96-well plates for spectrophotometric measurement at 560 nm. For epithelial-mesenchymal transition (EMT) assay, the expression of vimentin and E-cadherin was performed by western blot analysis.

### 2.7 Cell cycle and sub-G1 assay

The treated CT26 cells were collected, washed with PBS, fixed, and permeated with ice-cold 70% ethanol overnight, followed by incubation with 0.1% Triton X-100, 0.2 mg/mL RNaseA (Sigma) and staining with 20 μg/mL propidium iodide (PI) for 30 min at 37°C in the dark, and then analyzed by flow cytometry within 1 h. To determine the DNA histogram of cells, data acquisition and analysis were performed on a FACScan flow cytometer with CellQuest software (BD Biosciences, Franklin Lakes, NJ, USA).

### 2.8 Prediction and western blot assay of regulatory pathways

The targeted genes regulated by miR-762 were predicted by the MIRDB (http://mirdb.org/miRDB/policy.html) and TargetScan systems (http://www.targetscan.org/mmu_61/), while regulatory pathways were analyzed using DAVID Bioinformatics Resources software (NIAID/NIH, http://david.abcc.nciferf.gov/home.jsp). For western blot analysis of predicted molecule expression, the treated CT26 cells were cultured for 24 h, scraped cells from the culture dish, disrupted in 2× concentrated electrophoresis sample buffer (1 M Tris, pH6.8, 5% SDS, 40% glycerol, 0.005% bromophenol blue, and 8% β-mercaptoethanol), centrifuged for 5 min to pellet insoluble material, and subjected to gel electrophoresis on 10% (wt/vol) SDS-polyacrylamide gels. Protein samples were transferred onto polyvinylidene fluoride (PVDF) membranes. Primary antibodies, including vimentin, E-cadherin, Wnt-1, β-catenin, and MMP-9, were added to the PVDF membrane in TBST (50 mM Tris–HCl, pH 7.4, 150 mM NaCl, 0.1% Tween-20) containing 5% non-fat milk overnight at 4°C. Subsequently, the PVDF membrane was washed with TBST and incubated with appropriate secondary antibodies (horseradish peroxidase-conjugated goat anti-mouse or anti-rabbit IgG), followed by enhanced chemiluminescence (ECL; Visual protein, Taipei, Taiwan) and the membrane was examined by ECL spectrophotometry (Perkin Elmer, Geliance 600 Camera Imaging system, Waltham, MA, USA). GAPDH was used as an internal control.

### 2.9 Patients and data collection

The protocols for obtaining clinical samples and information were approved by the Institutional Review Board of National Taiwan University Hospital Yunlin Branch (No. 201611007RINB). Written informed consent to participate in this study was obtained from all of the individuals. This study collected blood samples from CRC patients (n = 20) and control donors (n = 20) (**Table 1**). CRC was diagnosed based on histological examination of surgical specimens and classified according to TNM staging (Union for International Cancer Control, UICC). Patients’ basic data were retrieved from electronic medical records and analyzed for age, gender, and tumor staging. All of the 20 CRC donors are adenocarcinoma (T2 – T4, M0 – M1), including colon (n = 14) and rectal (n = 6) cancers. Although CRCs typically start as a growth of tissue called a polyp, the present study included non-malignant symptom constipation (n = 1), colon polyp (n = 6), and hemorrhoid (n = 13) as control donors. The donors’ serum miRNAs were isolated with a TRIZol kit (https://www.ncbi.nlm.nih.gov/pmc/articles/PMC4793896/) and the levels of miR-762 were assessed by a real-time reverse transcription-polymerase chain reaction (RT-PCR) assay with a TaqMan™ MicroRNA Reverse Transcription Kit (https://www.ncbi.nlm.nih.gov/pmc/articles/PMC2225395/pdf/1746-4811-3-12.pdf). bdi-miR-159-3p RNA (10 ng/mL serum) was used as a control, and the primers are shown in **Table 2**.

**Table 1.** Characters of control donors and colorectal cancer (CRC) patients

| Characters |  | Control (n = 20) |  | CRC (n = 20) |  |
| --- | --- | --- | --- | --- | --- |
| Gender |  | Female: 6 | Male: 14 | Female: 6 | Male: 14 |
| Age<br>(year) | Mean ± SE | 63.5 ± 2.0 | 67.0 ± 2.7 | 57.2 ± 4.4 | 64.4 ± 4.3 |
|  | Range | 57–69 | 50–87 | 39–70 | 41–88 |
| Diagnosis |  | Constipation: |  | Colon cancer: |  |
|  |  | 0 | 1 | *T2-T3/M0:* |  |
|  |  |  |  | 3 | 6 |
|  |  | Colon polyp: |  | *T3-T4/M1:* |  |
|  |  | 0 | 6 | 0 | 5 |
|  |  | Hemorrhoid: |  | Rectal cancer: |  |
|  |  | 6 | 7 | *T2-T3/M0:* |  |
|  |  |  |  | 3 | 2 |
|  |  |  |  | *T3-T4/M1:* |  |
|  |  |  |  | 0 | 1 |

**Table 2.**
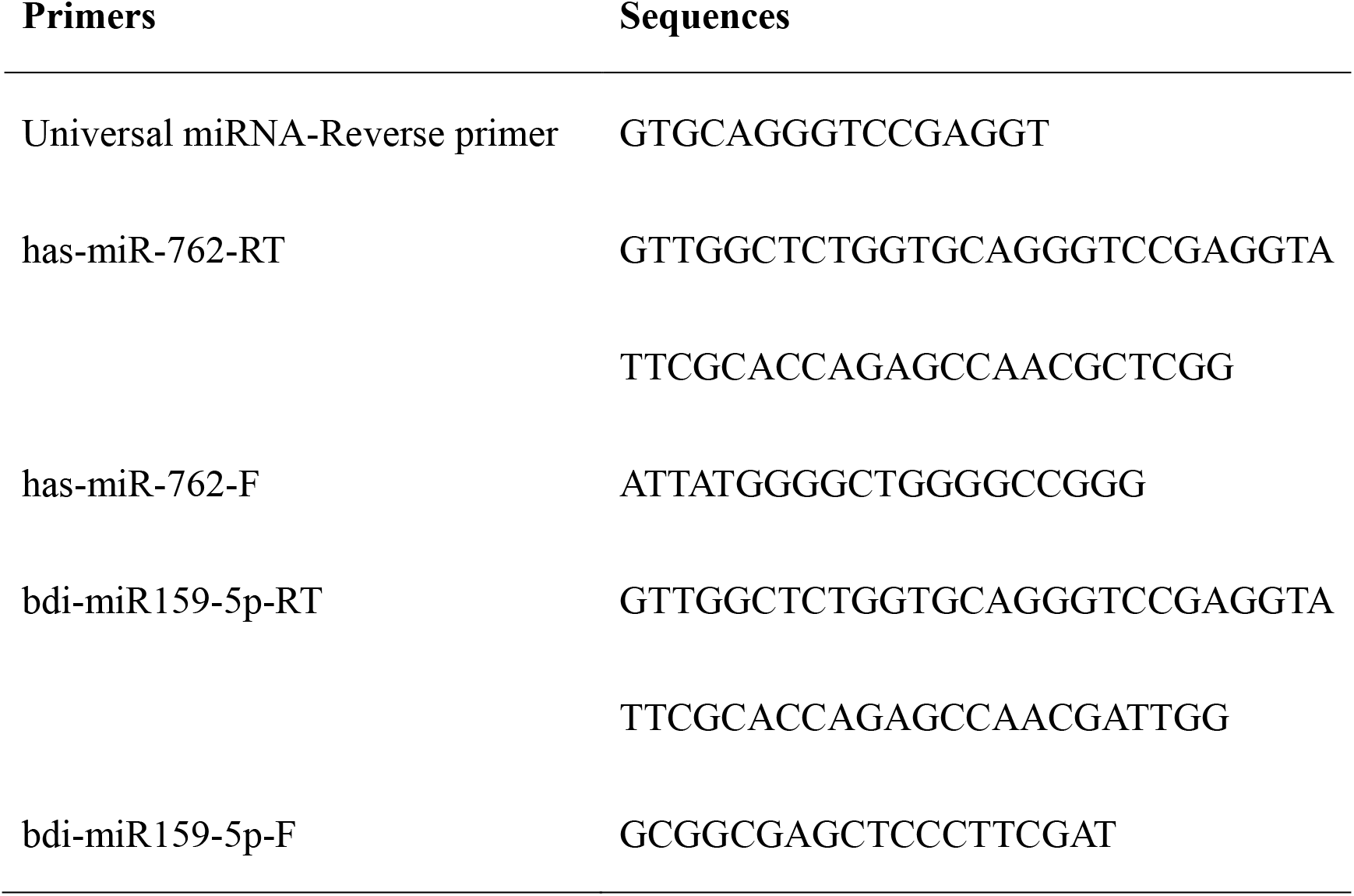
Primers used in this study

### 2.10 Statistical analysis

The results were expressed as the mean ± standard deviation (SD) of at least three experiments. Statistical comparisons were conducted with Student’s *t*-test or analysis of variance. A value of p < 0.05 was considered to indicate a statistically significant difference. All statistical analyses were performed using SigmaStat software (Systat Software, San Jose, CA, USA).

## 3 Results

### 3.1. Expression of mmu-miR-762 in BALB/c mice implanted with CRC CT26 cells

**Figure 1A** shows the animal experiment flow of BALB/c mice implanted with CRC CT26 cells, and plasma samples from each group were used for miRNA array analysis. Based on the animal experimental design, differentially expressed miRNAs for each comparison were determined by clustering analysis. Standard selection criteria for identifying differentially expressed miRNAs were established at log2|fold-changeļ ≥ 0.8 and p < 0.05. As shown in **Figure 1B**, plasma mmu-miR-762 levels in CT26 cell-implanted BALB/c mice were higher than those in normal mice according to miRNA array analysis, and the log2 ratio was 2.21.

### 3.2 Viability of CRC CT26 cells transfected with mmu-miR-762 mimic and inhibitor

As shown in **Figure 2A**, cells transfected with mmu-miR-762 mimic showed increased cell viability, whereas those transfected with mmu-miR-762 inhibitor showed decreased viability. However, transfection with the mimic and inhibitor of mmu-miR-762 did change the morphology of CT26 CRC cells (**Figure 2B**). CT26 cells transfected with mmu-miR-762 mimic and inhibitor also showed increased and decreased colony formation, respectively, suggesting that mmu-miR-762 upregulated cell proliferation (**Figure 2C**).

**Figure 2.**
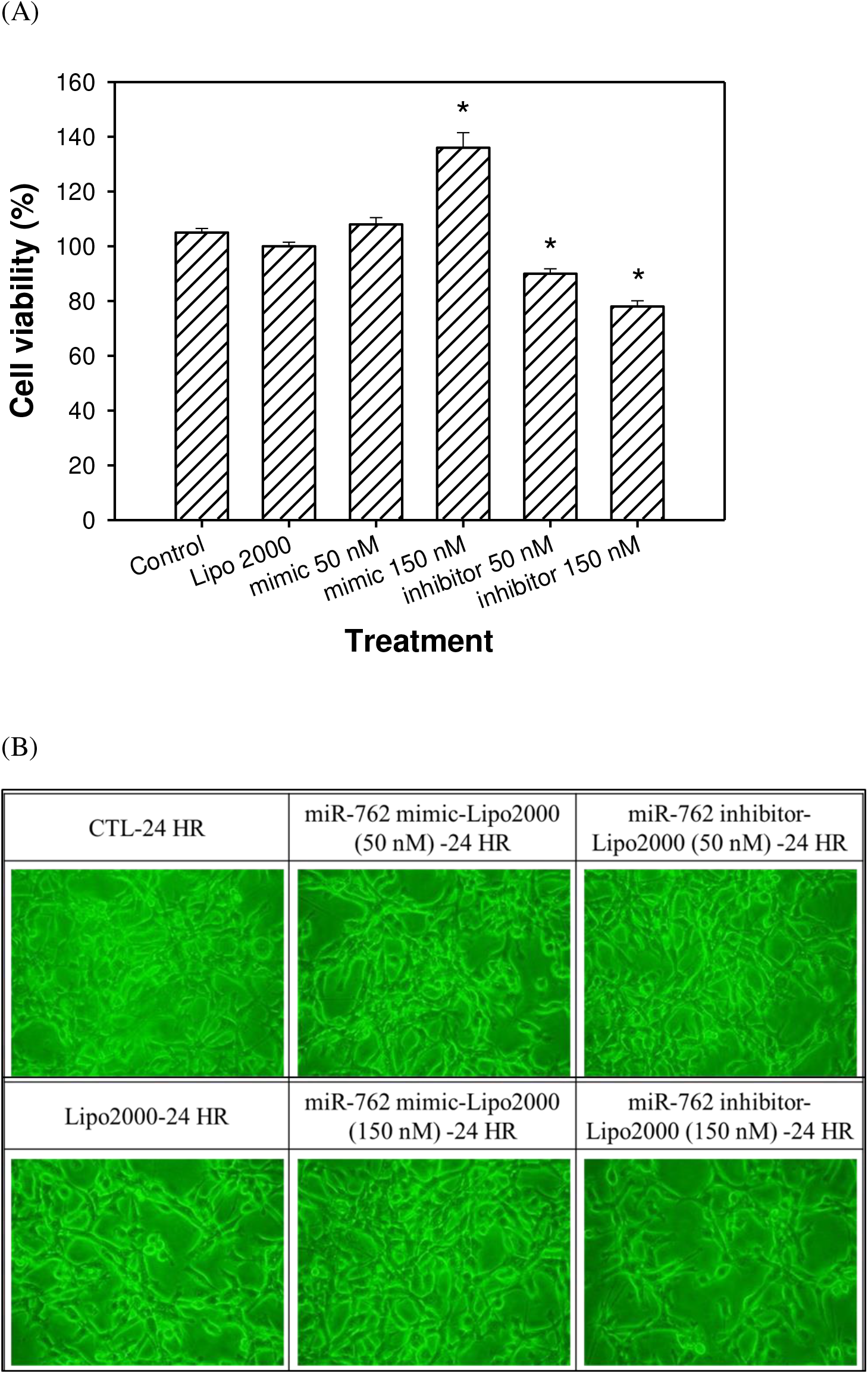
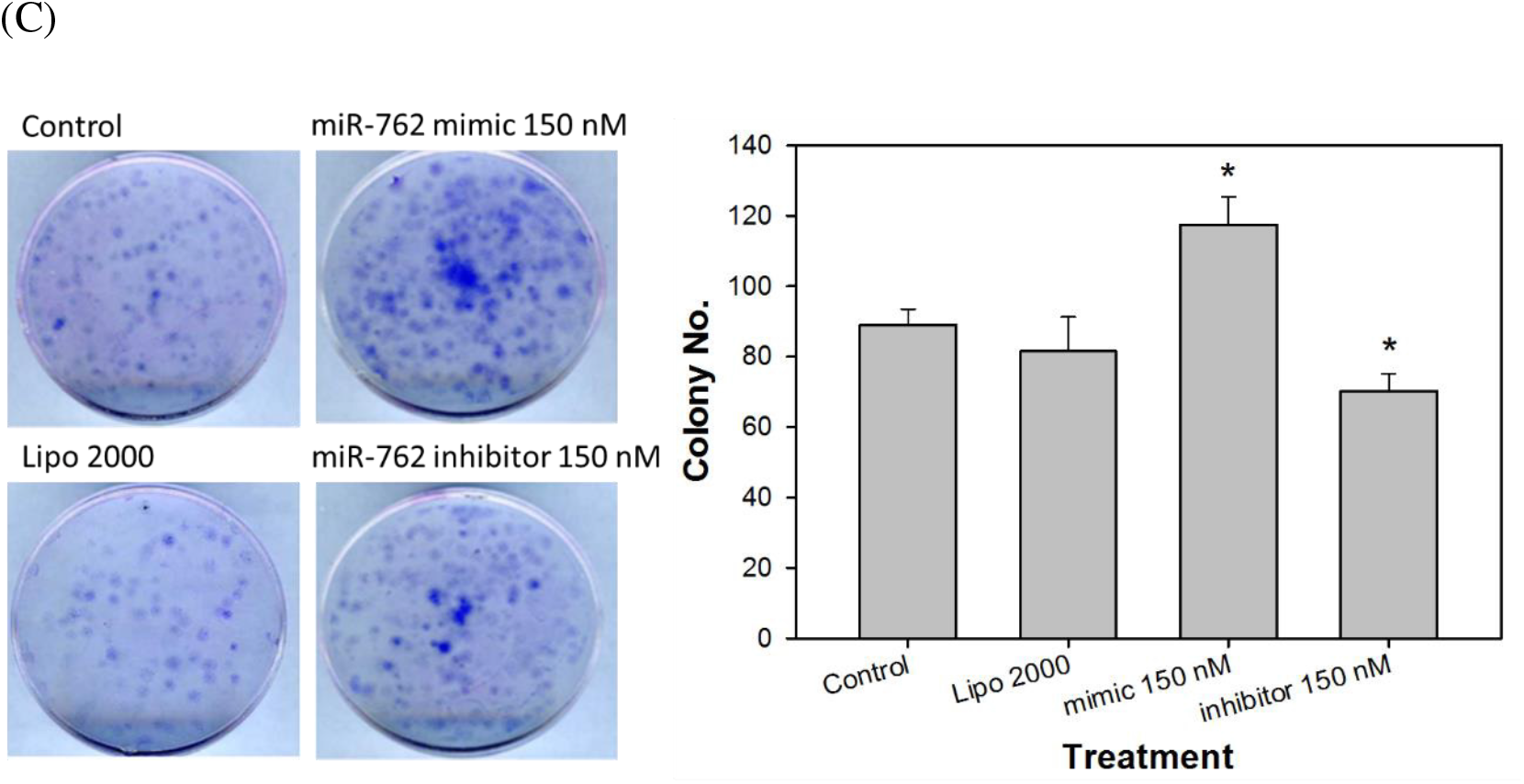
(A) Viability assay, (B) Morphological observation (200X), (C) Colony assay of CT26 cells transfected with mimic and inhibitor of mmu-miR-762. Cells transfected with mmu-miR-762 mimic (150 nM) increased cell viability and colony number, while mmu-miR-762 inhibitor (50 nM and 150 nM) decreased viability and colony formation by MTT assay. CT26 cells transfected with mimic and inhibitor of mmu-miR-762 did not show altered morphology under an inverted microscope view. The results were expressed as the mean ± standard deviation (SD) for three independent experiments. Statistical comparisons were conducted with Student’s *t*-test.* *P* < 0.05.

### 3.3 Invasion of CRC CT26 cells transfected with mmu-miR-762 mimic and inhibitor

As shown in **Figure 3A**, cells transfected with mmu-miR-762 mimic showed increased cell invasion ability, while the mmu-miR-762 inhibitor decreased cell invasion. This suggests that mmu-miR-762 regulates specific genes related to CRC cell invasion. Additionally, mmu-miR-762 mimic transfection to CT26 cells increased vimentin and decreased E-cadherin expression, suggesting that mmu-miR-762 might upregulate EMT (**Figure 3B**).

**Figure 3.**
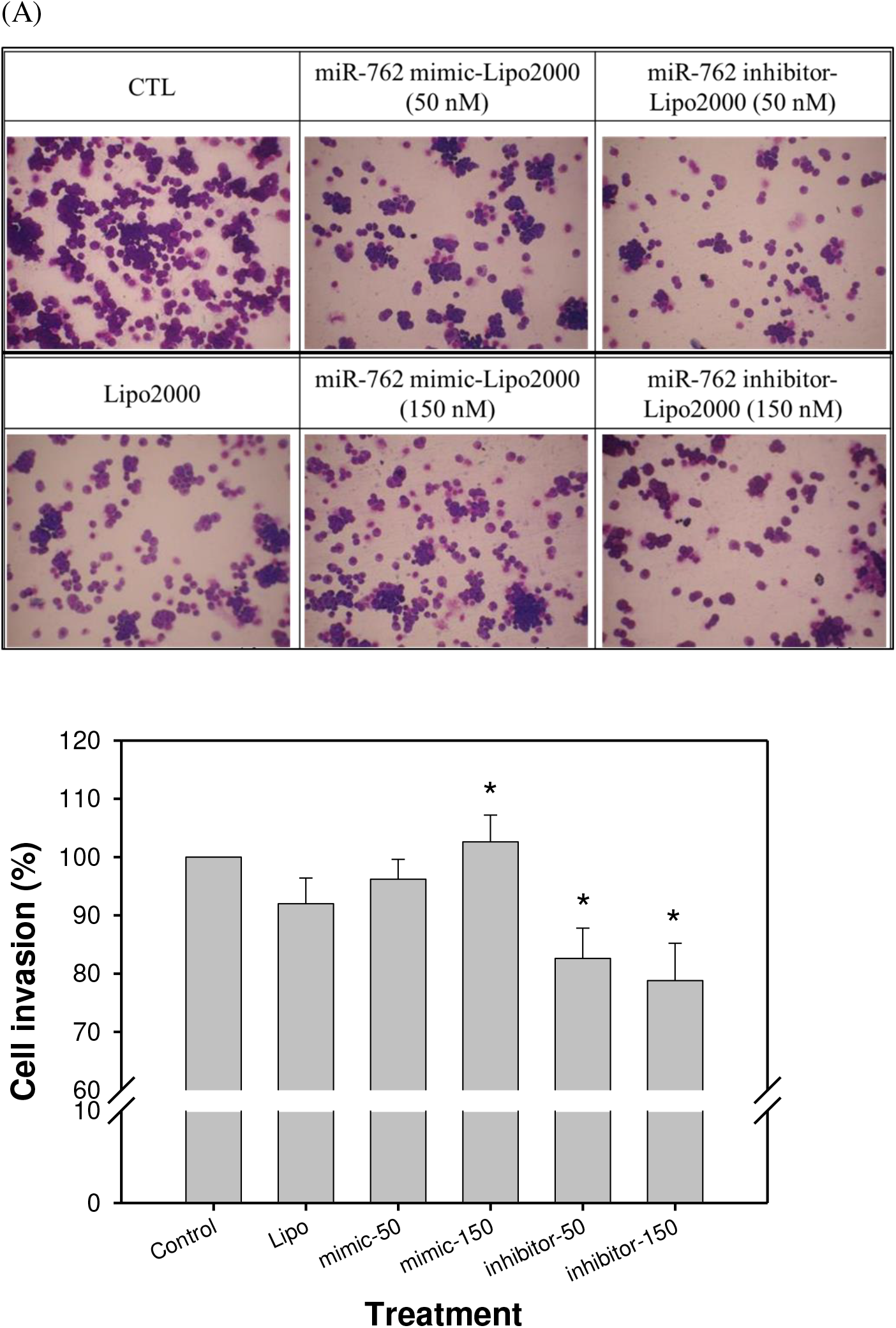
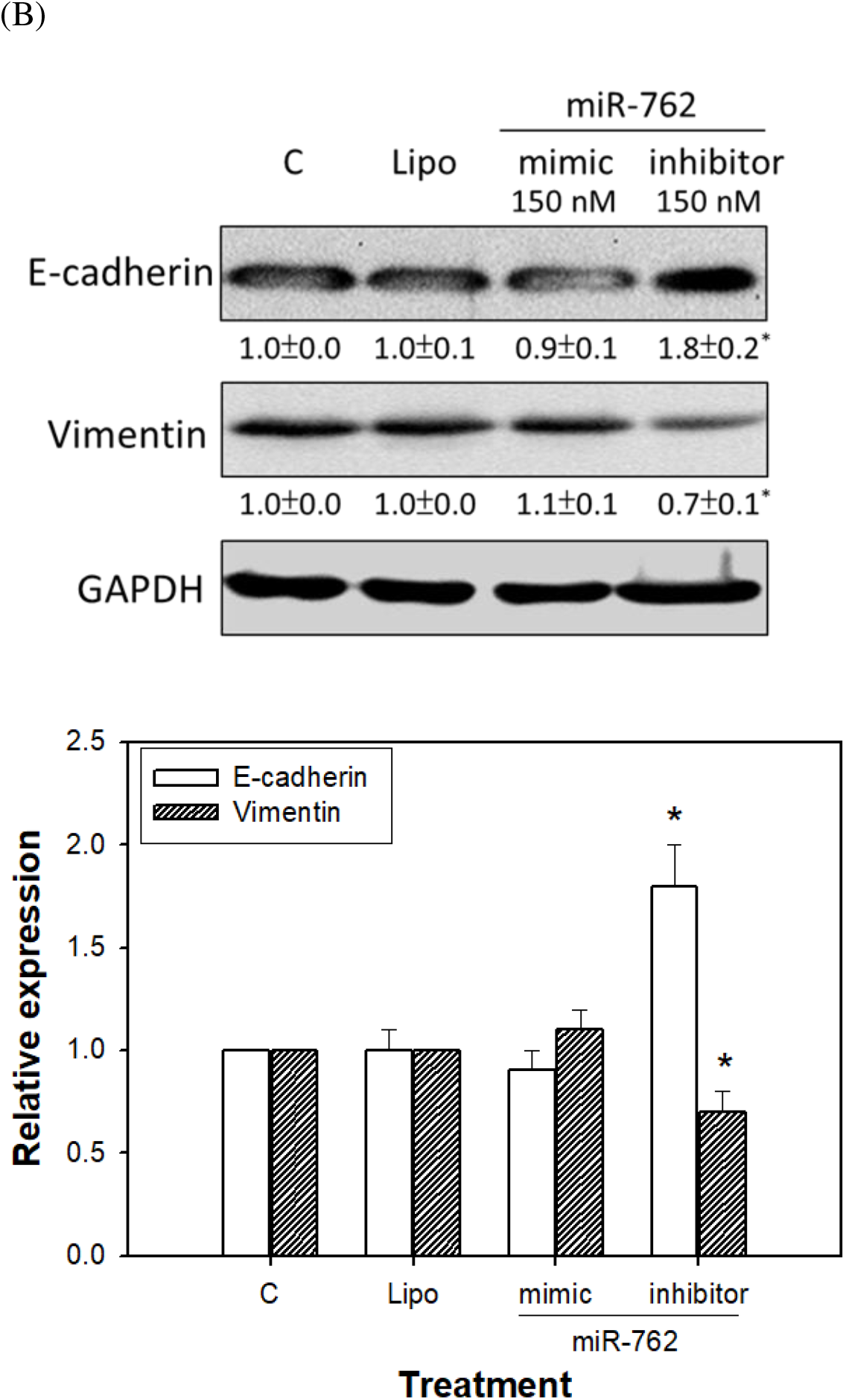
Invasion assay of CT26 cells transfected with mimic and inhibitor of mmu-miR-762. (A) Morphological observation of treated cells (100X), (B) Percent of invasive cells, (C) Expression of vimentin and E-cadherin by western blot analysis. Cells transfected with mmu-miR-762 mimic (150 nM) showed increased cell invasion ability, while mmu-miR-762 inhibitor (50 nM and 150 nM) decreased cell invasion. The results were expressed as the mean ± standard deviation (SD) for three independent experiments. Statistical comparisons were conducted with Student’s *t*-test. * *P* < 0.05.

### 3.4 Cell cycle of CRC CT26 cells transfected with mmu-miR-762 mimic and inhibitor

**Figure 4A** shows that both the mmu-miR-762 mimic and inhibitor did not change sub-G1 levels in CT26 cells, suggesting that mmu-miR-762 does not regulate cell apoptosis. Moreover, cells treated with high concentration (150 nM) of mmu-miR-762 inhibitor caused cell cycle arrest at G0/G1 phase (**Figure 4B**).

**Figure 4.**
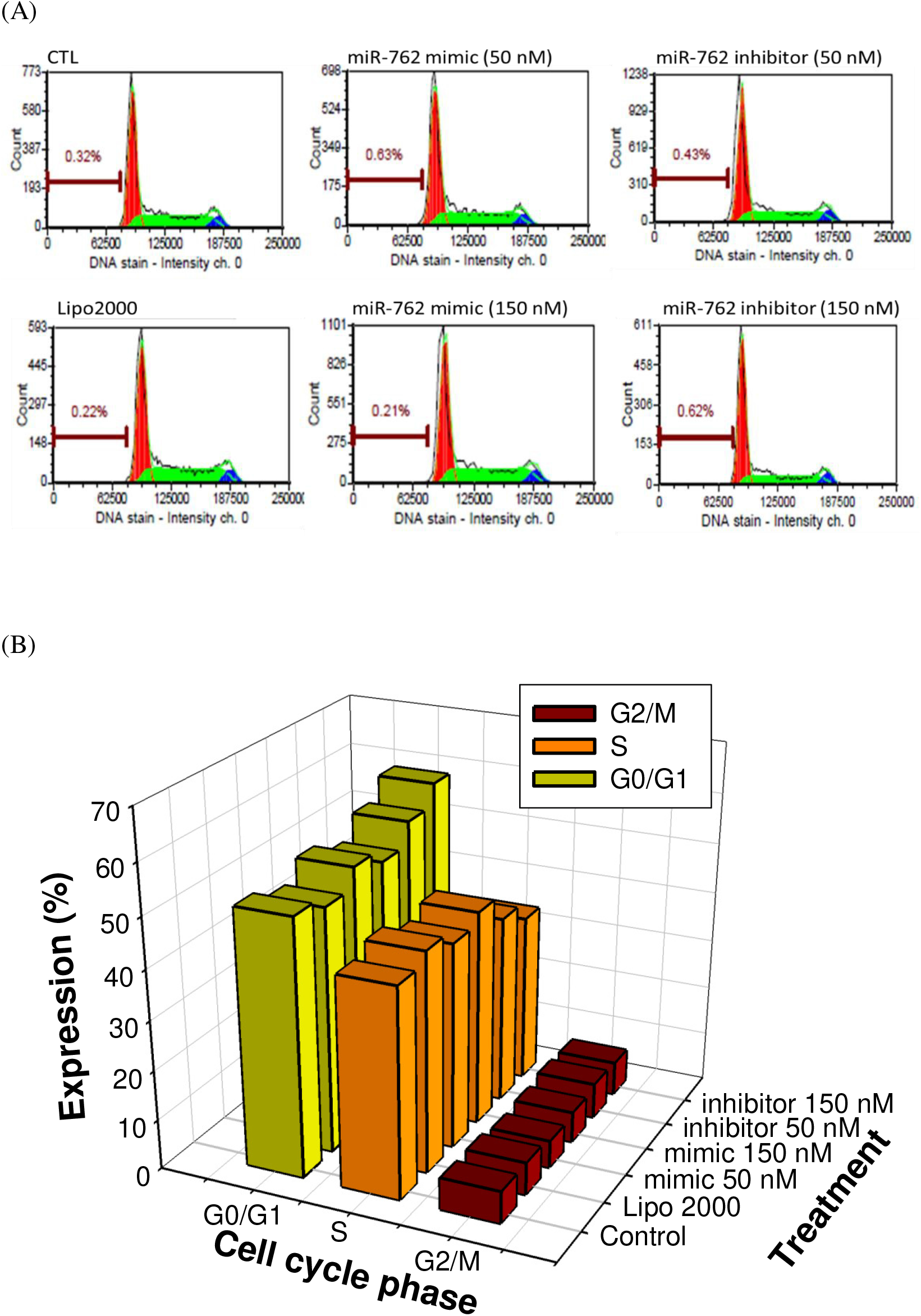
(A) Apoptosis sub-G1, (B) Cell cycle assay of CT26 cells transfected with mimic and inhibitor of miR-762. Neither mmu-miR-762 mimic nor inhibitor changed the sub-G1 levels of CT26 cells analyzed by a FACScan flow cytometer. The cell cycle arrest at G0/G1 phase when CT26 cells treated with 150 nM mmu-miR-762 inhibitor. The results were expressed as the mean ± standard deviation (SD) for three independent experiments. Statistical comparisons were conducted with Student’s *t*-test.

### 3.5 Signal molecule expression in CT26 cells transfected with mmu-miR-762 mimic and inhibitor

**Figure 5A** lists the predicted pathways that may be regulated by mmu-miR-762. The calcium signaling pathway, endocytosis, MAPK signaling pathway, and Wnt signaling pathway may be related to mmu-miR-762 regulation in CT26 cells.

**Figure 5.**
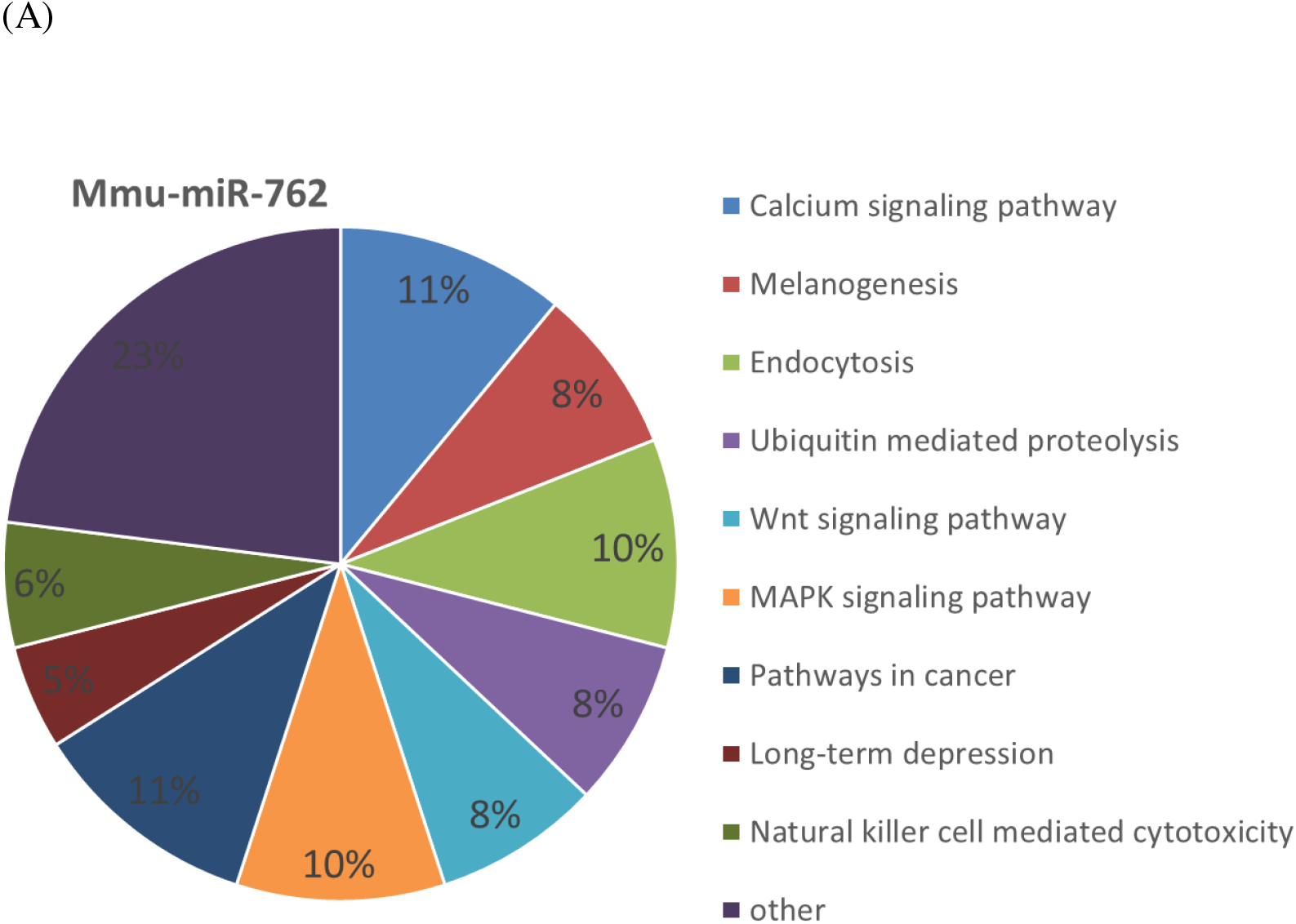
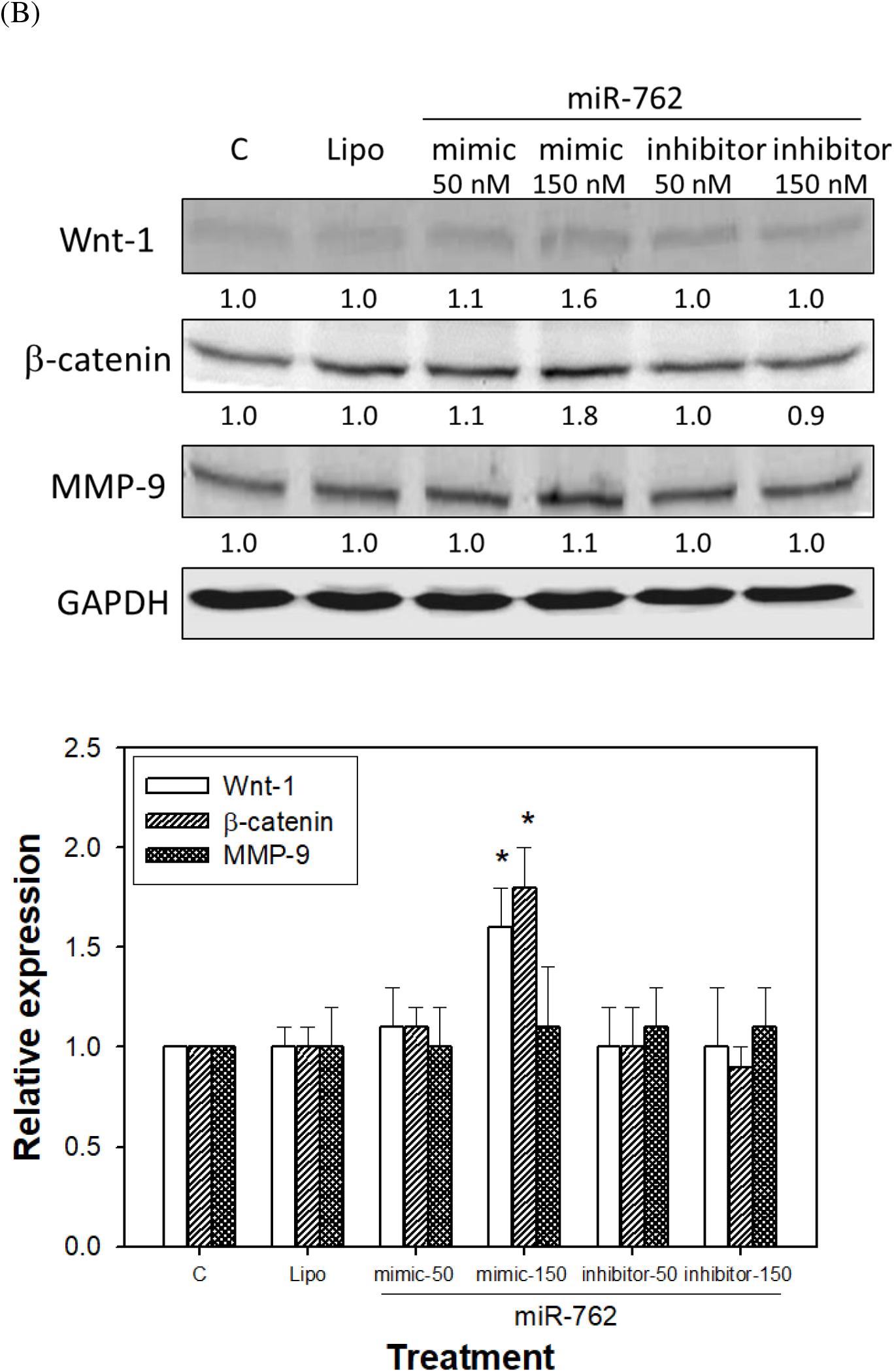
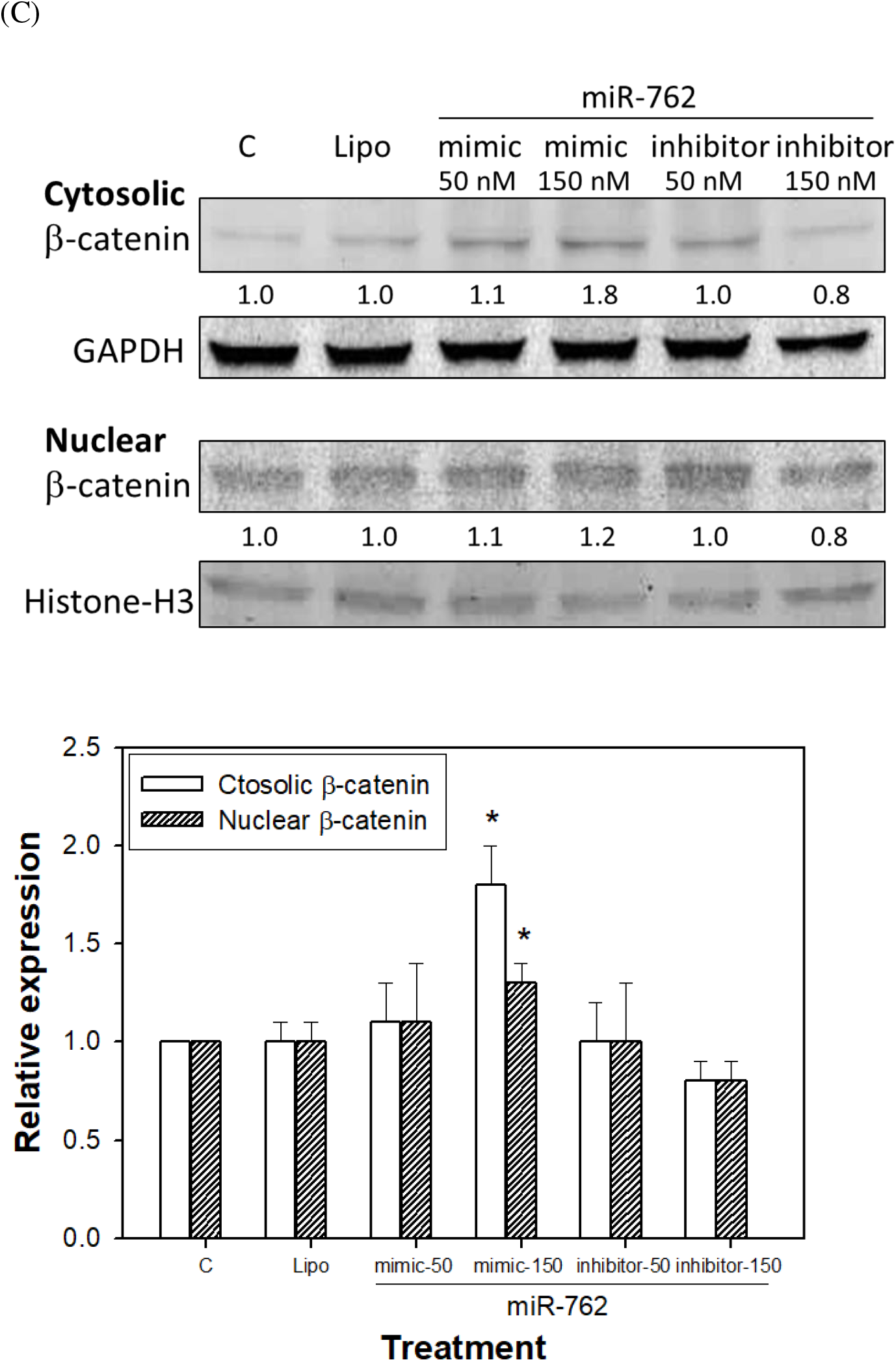
Wnt/β-catenin assay of CT26 cells transfected with mimic and inhibitor of miR-762. (A) Prediction of pathways of mmu-miR-762 assayed by DAVID Bioinformatics Resources software (NIAID/NIH), (B) Expression of Wnt-1, β-catenin, and MMP-9 by western blotting, and (C) Nuclear and cytosolic levels of β-catenin. The mmu-miR-762 mimic increased the expression of Wnt-1 and β-catenin, while mmu-miR-762 inhibitor decreased the levels of these molecules. MMP-9 was not changed in either mimic- or inhibitor-transfected cells. The cytosolic and nuclear levels of transcription factor β-catenin were both up-regulated in miR-762 mimic-transfected cells. The results were expressed as the mean ± standard deviation (SD) for three independent experiments. Statistical comparisons were conducted with Student’s *t*-test. * *P* < 0.05.

Western blot analysis showed that the mmu-miR-762 mimic increased the expression of Wnt-1 and β-catenin, while the mmu-miR-762 inhibitor decreased the expression of these molecules (**Figure 5B**). However, MMP-9 was not changed in either mimic- and inhibitor-transfected cells. Additionally, the cytosolic and nuclear levels of transcription factor β-catenin were both up-regulated in mmu-miR-762 mimic-transfected cells (**Figure 5C**), indicating that mmu-miR-762 induced the nuclear translocation of β-catenin. The results suggest that mmu-miR-762 regulated CRC CT26 cell viability and invasion, possibly through the Wnt/β-catenin pathway.

### 3.3 Serum has-miR-762 level in CRC cancer patients

The human subject solicitation were performed form April to October, 2017. **Table 1** shows the basic characters of control donors (n = 20) and CRC patients (n = 20). The CRC patients are diagnosed for the first time or relapsed, participating in this project before receiving clinical treatment. **Table 2** lists the primers used for the RT-PCR assay of serum has-miR-762 levels. There were no significant differences in gender and age between the two groups. The serum has-miR-762 level in CRC cancer patients was significant higher than in control donors (**Figure 6A**). Relative has-miR-762 expression in CRC patients (T2 – T4, M0 – M1) was 464.7 ± 142.3, while control donors showed a value of 53.2 ± 11.7 (p = 0.0073). According to the TNM staging, CRC patients with distant metastasis (M1, n = 6) showed significantly higher expression of serum has-miR-762 than CRC patients without distant metastasis (M0, n = 14) (M0 vs. M1, 129.2 ± 142.1 vs. 724.5 ± 872.3, p = 0.019) (**Figure 6B**). These results in animal, cell culture, and CRC patients suggested that has-miR-762 is a crucial factor that stimulates CRC growth and invasion, and such effects may be regulated by the Wnt/β-catenin pathway.

**Figure 6.**
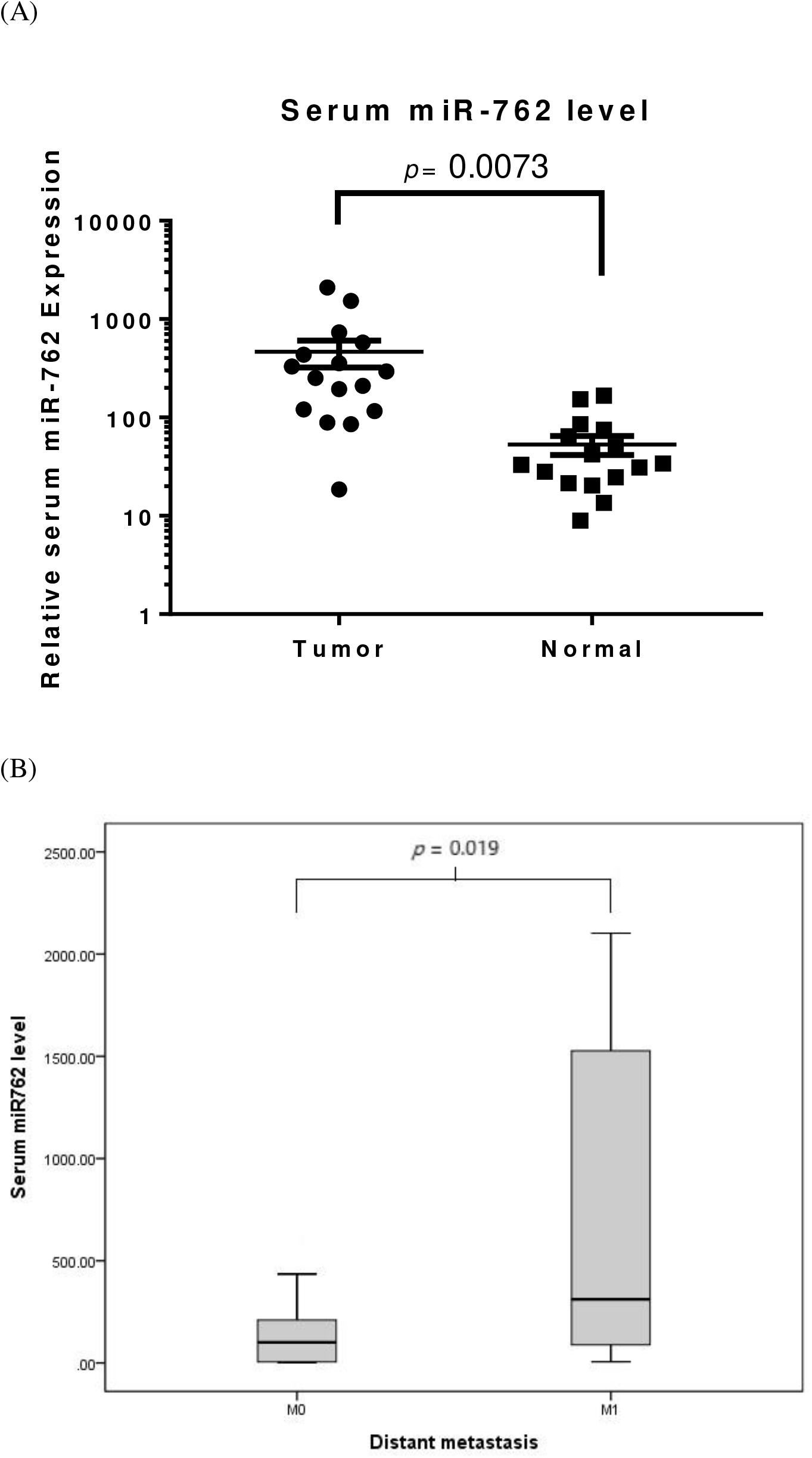
(A) Serum has-miR-762 expression in CRC patients and in control donors. Serum has-miR-762 expression in CRC patients (n = 20) was significantly higher than in control donors (n = 20), (B) The box plot shows the median of serum has-miR-762 expression, and the box shows the 25–75th percentile of serum has-miR-762 expression. CRC patients with distant metastasis (M1, n = 6) had higher expression of serum has-miR-762 than CRC patients with no distant metastasis (M0, n = 14). The results were expressed as the mean ± standard deviation (SD) for three independent experiments. Statistical comparisons were conducted with Student’s *t*-test. * *P* < 0.05.

## 4. Discussion

miRNAs are a group of endogenous, small (18–25-nucleotide long), and non-coding RNA molecules that regulate the expression of specific mRNAs by either translational inhibition or mRNA degradation^5^. More than 50% of miRNAs are reported to be in cancer-associated genomic break points and function as tumor suppressors or oncogenic molecules^6^. Among the cancers, most CRCs form from the normal mucosa through the adenoma stage, which is accompanied by numerous gene mutations. Our study has screened circulating mmu-miR-762 expression in mice with CRC CT26 cell implantation. Further study demonstrate that CT26 cells transfected with mmu-miR-762 mimic promoted the cell viability and invasion/EMT of CRC cells. A similar result was also seen in human subjects. Expression of mmu-miR-762 in circulation was significantly higher in clinical CRC patients than in control donors. We subsequently confirmed that mmu-miR-762 significantly increased in distant metastasis of CRC, resulting in poor prognosis. Our study of the biological roles of miR-762 in CRC development revealed Wnt/β-catenin as a downstream signaling pathway. These findings suggest that miR-762 plays a fundamental role in colorectal cancer tumorigenesis by affecting cancer cell proliferation and invasion/EMT.

miR-762 is upregulated in radiation-induced tumors in mice and may affect the pathways involved in apoptosis in this context^15^. Previous studies reported abnormal miRNA expression in numerous cancers, and miR-762 overexpression has been reported in breast cancer and ovarian cancer, where it promoted cancer cell growth and metastasis^16,17^. A previous study also demonstrated an association between miR-762 expression and oral carcinogenesis^18^. However, no studies have examined the expression and function of circulating miR-762 in CRC. In this study, CRC cell line with mmu-miR-762 transfection was frequently up-regulated the cell viability and EMT, as well as has-miR-762 detected in serum extracted from CRC patients. Overexpression of mmu-miR-762 enhanced CRC cell proliferation and invasion. The ability of miR-762 to promote cell proliferation and invasion was confirmed by both overexpression and down-regulation experiments *in vitro.* Therefore, our results suggest that miR-762 is a novel oncogenic RNA in CRC.

To explore the molecular mechanisms by which miR-762 enhances CRC cell growth and invasion, Wnt/β-catenin was identified as a direct target of mmu-miR-762 in CRC cells. First, the Wnt/β-catenin signaling pathway was predicted to be regulated by miR-762^19,20^. Second, western blot analysis showed that the miR-762 mimic increased the expression of Wnt-1 and β-catenin. Third, the cytosolic and nuclear levels of transcription factor β-catenin were both up-regulated in miR-762 mimic-transfected cells. However, CT26 cells transfected with the mimic and inhibitor of miR-762 did not show altered sub-G1 levels in CT26 cells. The cell cycle was arrested at G0/G1 phase in cells treated with high concentration (150 nM) of miR-762 inhibitor. Additionally, MMP-9 was not changed in either mimic- and inhibitor-transfected cells. These data suggest that mmu-miR-762 regulates CRC CT26 cell viability and invasion/EMT might through the Wnt/β-catenin pathway, but does not regulate cell cycle and apoptosis.

Although miR-762 is overexpressed in several cancers, no studies have investigated miR-762 in CRC. In this study, real-time PCR analysis of human subjects confirmed that the expression of serum has-miR-762 in CRC patients was much higher than that in control donors. There was also an association between serum has-miR-762 expression and CRC patients with distant metastasis. miR-762 exerted its regulatory effects on cell proliferation and invasion/EMT by affecting the Wnt/β-catenin signaling pathway. β-Catenin can influence metastasis through its effects on the EMT^21^. An article reported that FOXP3 interacts with β-catenin could induce transcription of Wnt target genes to promote cell proliferation, invasion and EMT induction^22^. The present study demonstrated a similar result in CRC that miR-762 promoted CRC cell metastasis by promoting the Wnt/β-catenin pathway according to the results of the Transwell invasion assay and EMT molecules (vimentin and E-cadherin) by Western blotting.

To confirm that human miR-762 can regulate Wnt pathway as well as mouse miR-762, we used the TargetScan microRNA target prediction which algorithm is considered the conservation among species. The target prediction results were listed of both human hsa-miR-762 target (supplementary table S1) and murine mmu-miR-762 target prediction (supplementary table S2), demonstrating that the Wnt ligands were regulated by miR-762 in both human and murine. In human, has-miR-762 has predicted to suppress Wnt family such as 7B, 2B, 3A, 9A, 5A, 10B, 3, 7A, and 11. In mmu-miR-762, it has predicted to suppress Wnt ligands 9B, 7B, 6, 10B, 10A, 4, 3A, 7A, and 8. Among these, Wnt-7B, −3A, and −10B have the genetic conservation of miR-762 binding site between human and murine 3’-UTR. Although the miR-762 has certain different target specify among human and mice, they still share some intersection in Wnt signaling. In the future, further analysis of the regulation of circulating miR-762 on cancer promotion and progression in CRC patients will be advanced elucidated.

Hou et al. (2017) reported that miR-762 can downregulate the expression of a tumor suppressor protein menin through a binding site in its 3’-UTR and consequently upregulate the Wnt cell signaling pathway to promote the development of ovarian cancer^17^. Our present study first revealed that circulating miR-762 expression may be useful as a biomarker for diagnosing CRC and higher serum miR-762 expression-caused distant metastasis of patients with CRC. While patients with high expression of miR-762 may have shorter survival times than those with low expression of miR-762, survival analysis could not be performed because of the relatively small sample size and short time. Furthermore, the associations between serum miR-762 and tumor size and lymph node invasion were not thoroughly investigated. Therefore, further large-scale studies are necessary to investigate whether serum miR-762 levels can be used as a predictive and a prognostic factor of patients with CRC.

## 5. Conclusions

The present study demonstrated the following. (1) In BALB/c mice, plasma mmu-miR-762 levels in CT26 cell-implanted mice were higher than those in normal mice determined by miRNA array analysis. (2) In CRC CT26 cells, transfection of mmu-miR-762 inhibitor decreased cell viability, colony formation, cell invasion, EMT, and Wnt/β-catenin expression. (3) In humans, serum has-miR-762 levels in CRC patients were higher than in control donors according to the RT-PCR assay. (4) Higher serum has-miR-762 levels in CRC patients induced distant metastasis. Therefore, we suggested that miR-762 upregulation in CRC may be useful as a biomarker for the clinical diagnosis and prognosis. New drug development of miR-762 inhibitors or blockers may be an effective strategy for CRC treatment.

## Abbreviations

ATCC: American Type Culture Collection
CO2: carbon dioxide
CRC: colorectal cancer
DHHS: Department of Health and Human Services
DNA: deoxyribonucleic acid
ECL: enhanced chemiluminescence
EDTA: ethylenediaminetetraacetic acid
EMT: epithelial-mesenchymal transition
FBS: fetal bovine serum
GAPDH: glyceraldehyde 3-phosphate dehydrogenase
IACUC: institute animal care and use committee
miR, miRNA: microRNAs
MAPK: mitogen-activated protein kinase
MIRDB: microRNA target prediction database
MMP-9: matrix metallopeptidase 9
MTT: 3-(4,5-Dimethylthiazol-2-yl)-2,5-diphenyltetrazolium bromide
NIH: National Institutes of Health
PBS: phosphate-buffered saline
PI: propidium iodide
PVDF: polyvinylidene fluoride
RPMI: Roswell Park Memorial Institute
RT-PCR: real-time reverse transcription-polymerase chain reaction
SE: standard error
SD: standard deviation
SDS: sodium dodecyl sulfate
TBST: Tris buffered saline with Tween-20
UICC: Union for International Cancer Control

## Author Contribution

Liao HF and Chen CY conceived and designed the experiments, Chang WM, Chen YY and Lin YF performed the experiments, Lai PS, Liao HF and Chen CY analyzed the data and wrote the paper.

## Availability of data and materials

All materials are available from the corresponding author.

## Consent for publication

Not applicable

## Ethics approval and consent to participate

This study was approved by the Institutional Review Board of National Taiwan University Hospital Yunlin Branch (No. 201611007RINB).

## Acknowledgements

Nil

